# Constitutive activation of TORC1 signaling attenuates virulence in the cross-kingdom fungal pathogen *Fusarium oxysporum*

**DOI:** 10.1101/2022.08.08.503127

**Authors:** Gesabel Yaneth Navarro-Velasco, Antonio Di Pietro, Manuel Sánchez López-Berges

## Abstract

The filamentous fungus *Fusarium oxysporum* causes vascular wilt disease in a wide range of plant species and opportunistic infections in humans. Previous work suggested that invasive growth in this pathogen is controlled by environmental cues such as pH and nutrient status. Here we investigated the role of Target Of Rapamycin Complex 1 (TORC1), a global regulator of eukaryotic cell growth and development. Inactivation of the negative regulator Tuberous Sclerosis Complex 2 (Tsc2), but not constitutive activation of the positive regulator Gtr1, in *F. oxysporum* resulted in inappropriate activation of TORC1 signaling under nutrient limiting conditions. The *tsc2*Δ mutants showed reduced colony growth on minimal medium with different nitrogen sources and increased sensitivity to cell wall or high temperature stress. Furthermore, these mutants were impaired in invasive hyphal growth across cellophane membranes and exhibited a marked decrease in virulence, both on tomato plants and on the invertebrate animal host *Galleria mellonella*. Importantly, invasive hyphal growth in *tsc2*Δ strains was rescued by rapamycin-mediated inhibition of TORC1. Collectively, these results reveal a key role of TORC1 signaling in development and pathogenicity of *F. oxysporum* and suggest new potential targets for controlling fungal infections.

**AUTHOR SUMMARY:** Filamentous fungal pathogens cause devastating losses in agriculture and lethal infections in humans. A prerequisite of fungal infection is invasive hyphal growth, a process that is exquisitely controlled by environmental cues including nutrients and host signals. Here we examined the role of TOR complex 1 (TORC1), a conserved pathway that regulates cell growth in response to nutrient status. We show that deregulation of TORC1 signaling impairs pathogenicity in *Fusarium oxysporum*, a cross-kingdom fungal pathogen that attacks more than 150 different crops as well as immunocompromised humans. Inactivation of Tsc2, a negative regulator of TORC1, led to constitutive TORC1 activation, reduced growth under nutrient-limiting conditions and increased sensitivity to cell wall stress. Importantly, *tsc2*Δ mutants were impaired in invasive hyphal growth and in virulence on plant and animal hosts. Our results support a conserved role of TORC1 as a negative regulator of pathogenicity-related functions and reveal new leads for antifungal drug discovery.

## INTRODUCTION

Soil-borne fungal plant pathogens are ubiquitous and highly persistent, causing massive losses in field and greenhouse crops. Fungicide application, resistance breeding or crop rotation are agricultural practices that have proven insufficient to prevent root diseases [1]. The soil-inhabiting ascomycete *Fusarium oxysporum* causes vascular wilt in more than 150 different plant species and ranks among the most important phytopathogens [2]. Furthermore, this fungus can also produce opportunistic infections in humans ranging from superficial or locally invasive to disseminated depending on the immune status of the patient [3]. Strikingly, a single *F*. *oxysporum* f. sp. *lycopersici* isolate can infect and kill both tomato plants and immunosuppressed mice [4] making this an excellent model to study the genetic basis of cross-kingdom pathogenesis in fungi.

Previous studies established that plant infection by *F*. *oxysporum* is regulated by a network of conserved signaling pathways that sense and transduce environmental cues such as nutrients, ambient pH or host compounds [5–9]. One of the key pathogenicity mechanisms, invasive hyphal growth, was found to be dependent on nutrient status and promoted by rapamycin, a specific inhibitor of the protein kinase TOR (Target Of Rapamycin) [7]. TOR is broadly conserved in eukaryotes and controls nutrient signaling [10] and cell growth from yeast to mammals [11–13]. TOR interacts with other proteins to form two structurally and functionally distinct complexes, TOR complex 1 (TORC1) and 2 (TORC2) [11–13]. The rapamycin-sensitive TORC1 promotes cell growth in response to nutrients, growth factors and cellular energy by activating anabolic processes such as ribosome biogenesis and protein synthesis and preventing catabolic processes such as autophagy and ubiquitin-mediated proteolysis [14], while TORC2 is insensitive to rapamycin and controls actin cytoskeleton organization [15]. Activity of TORC1 is positively and negatively regulated, respectively, by the small GTPase Rheb [16] and its GTPase-activating protein (GAP) Tsc2 [15]. A second upstream component is the vacuolar membrane associated EGO protein complex [11, 17], with the small GTPases Gtr1/Gtr2 acting as TORC1 activators [17].

Currently, the role of TORC1 in fungal pathogenicity on plants is controversial. Genetic or pharmacological inhibition of TORC1 does affects virulence in different phytopathogens such as *Botrytis cinerea*, *Verticillium dahliae*, *Fusarium graminearum* and *F. oxysporum*, but this could be explained by a severe reduction of mycelial growth [18–21]. On the contrary, studies in the rice blast fungus *Magnaporthe oryzae* point towards an inhibitory function of TORC1 in appressorium formation and other virulence-related processes such as autophagy [22, 23]. In line with this, we previously found that pharmacological inhibition of TORC1 by rapamycin promotes invasive growth in *F. oxysporum*, suggesting that TORC1 negatively regulates pathogenicity functions [7].

Here we investigated the effect of constitutive TORC1 activation, through targeted deletion of *tsc2* (*tsc2*Δ) and/or expression of a constitutively active *gtr1^Q86L^* allele (*gtr1^GTP^*), on development and pathogenesis of *F. oxysporum*. We found that inappropriate activation of TORC1 in *tsc2*Δ strains leads to impaired colony growth under nutrient-limiting conditions, increased sensitivity to stresses, impaired invasive growth and reduced virulence on plant and animal hosts, highlighting the crucial role of TORC1 regulation during fungal development and pathogenicity.

## RESULTS

### Establishment of TORC1 readouts in *Fusarium oxysporum*

To measure TORC1 activity in *F. oxysporum*, we first tested the feasibility of two different readouts used previously in *Saccharomyces cerevisiae* and *Schizosaccharomyces pombe*: transcriptional activation of ribosomal genes [24, 25] and phosphorylation of the Ribosomal protein S6 (Rps6) [26, 27]. For the first readout we monitored transcript levels of the *F. oxysporum* ribosomal genes *FOXG_01615*, *FOXG_07725* and *FOXG_09465*, encoding 50S ribosomal protein L4e, 40S ribosomal protein S5 and 40S ribosomal protein S3, respectively, while for the second readout we determined the phosphorylation status of *F. oxysporum* Rps6 using a phospho-Akt antibody (see materials and methods).

In a short-term adaptation experiment, *F. oxysporum* germlings were either transferred from minimal medium lacking a nitrogen source (−N) to minimal medium with glutamine (Gln), or maintained in −N conditions. A shift from −N to Gln resulted in rapid upregulation of the three ribosomal genes and a marked increase in Rps6 phosphorylation, both indicative of TORC1 activation (Fig 1A). By contrast, the inverse shift from Gln to −N caused a decrease in both readouts, that was further exacerbated in the presence of 100 ng ml^-1^ rapamycin (−N+R) (Fig 1B). Importantly, the same effect of rapamycin treatment was observed even when the fungus was maintained in Gln, confirming the inhibitory effect of this drug on TORC1 activity (Fig 1C). We conclude that levels of ribosomal gene transcripts and of Rps6 phosphorylation can be used to reliably monitor TORC1 activity in *F. oxysporum*.

**Fig 1.**
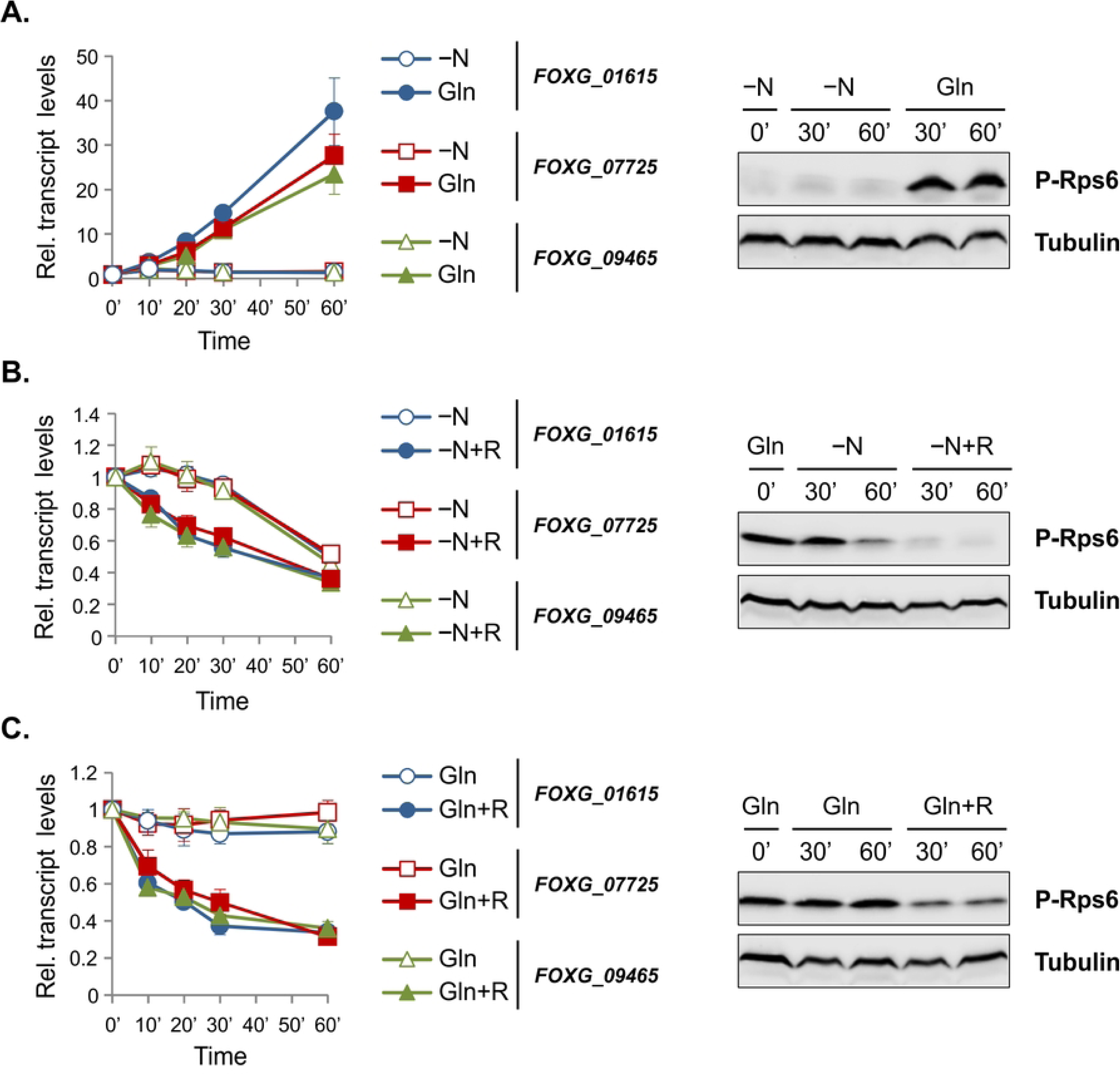
Establishment of TORC1 readouts in *Fusarium oxysporum*. (A-C) The wild-type strain was germinated 16 h in Potato Dextrose Broth (PDB) and then transferred to: (A) nitrogen-free minimal medium (−N) for 60 min (time point 0’) before shifting either to −N (empty symbols) or glutamine minimal medium (Gln) (solid symbols) for an additional 60 min; (B) Gln for 60 min (time point 0’) before shifting either to −N (empty symbols) or −N + 100 ng ml^-1^ rapamycin (-N+R) (solid symbols) for an additional 60 min; or (C) Gln for 60 min (time point 0’) before shifting to Gln (empty symbols) or Gln + 100 ng ml^-1^ rapamycin (Gln+R) (solid symbols) for an additional 60 min. *Left panels:* Transcript levels of the indicated ribosomal genes were measured at the indicated time points by quantitative real-time RT-PCR, normalized to the *act1* gene and expressed relative to those obtained at time 0’. Bars represent standard deviations calculated from two independent experiments with three technical replicates each. *Right panels:* Representative western blots showing phosphorylation of the ribosomal protein S6 (Rps6) at the indicated time points. α-Tubulin was used as loading control. Blots were performed with two biological replicates with similar results.

### Targeted deletion of *tsc2*, but not expression of a constitutively active *gtr1^Q86L^* allele (*gtr1^GTP^*), activates TORC1 signaling in *Fusarium oxysporum*

We attempted two different approaches for constitutive TORC1 activation in *F. oxysporum*: (1) deletion of *tsc2* (*tsc2*Δ) and (2) expression of a constitutively active *gtr1^Q86L^* allele (*gtr1^GTP^*). A BlastP search in FungiDB [28] using Tsc2 from *S. pombe* (SPAC630.13c) as a bait detected a single *F. oxysporum* Tsc2 orthologue (FOXG_01632). ClustalW alignment [29] of the GTPase-activating domain of Tsc2 from evolutionarily distant organisms showed a high degree of conservation (S1 Fig). Isogenic *tsc2*Δ mutants of *F. oxysporum* were obtained by replacing the entire *FOXG_01632* coding sequence with the *hygromycin B* resistance gene (S2 Fig). When the *tsc2*Δ mutant was submitted to a shift from Gln to −N, both ribosomal gene transcript and Rps6 phosphorylation levels remained largely stable, in contrast to the wild-type strain, suggesting that activation of TORC1 in the *tsc2*Δ mutant is independent of nutrient status (Fig 2A). As expected, rapamycin-mediated TORC1 inhibition still caused a decrease of TORC1 readouts in *tsc2*Δ, consistent with pathway hyperactivation occurring upstream of TORC1 (Fig 2A). Importantly, reintroduction of the wild-type *tsc2* allele into *tsc2*Δ, yielding the complemented strain *tsc2*Δ^C^ (S2 Fig), fully restored wild-type values for the TORC1 readouts upon a shift from Gln to −N (Fig 2B).

**Fig 2.**
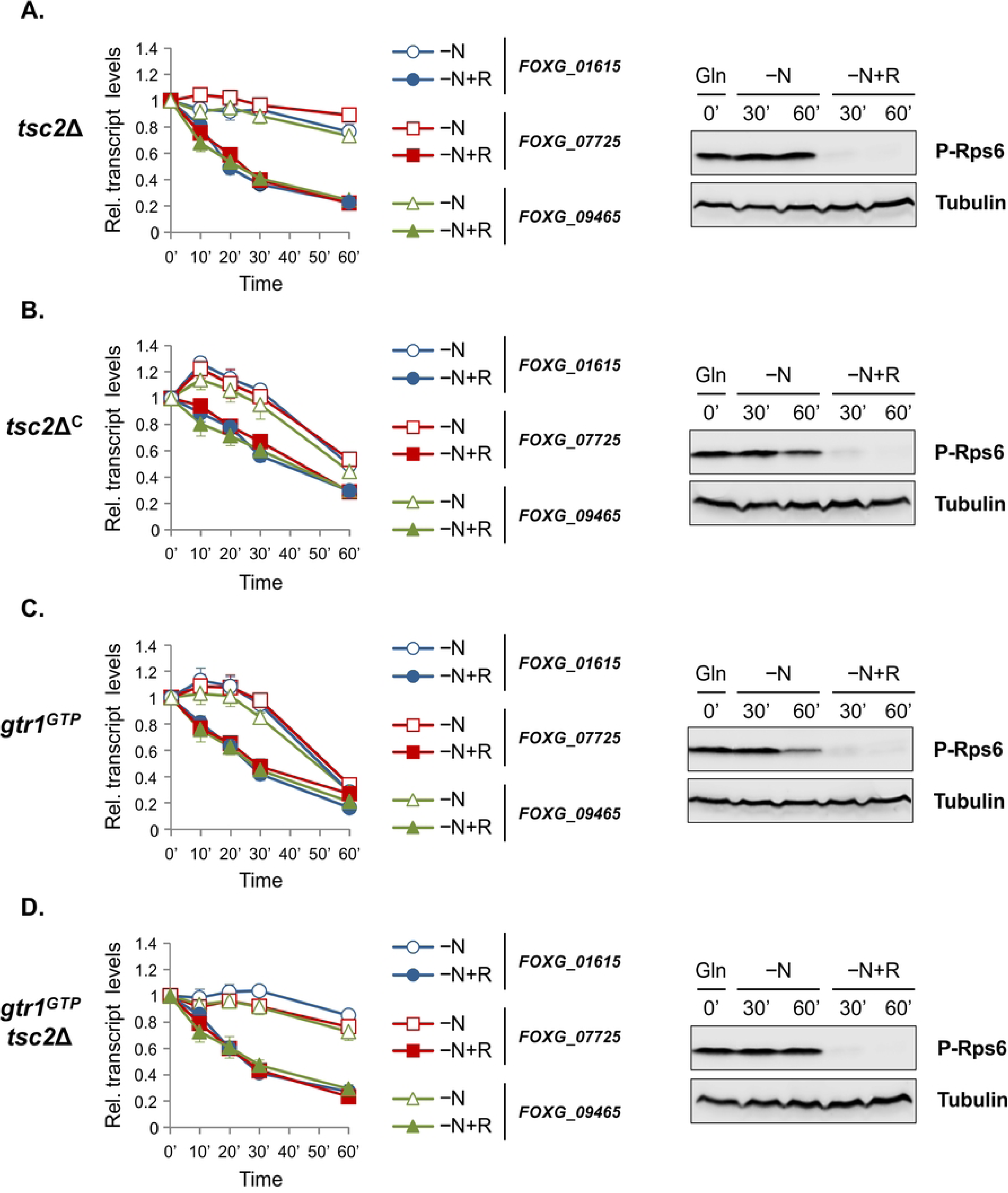
Targeted deletion of *tsc2* leads to nitrogen status-independent TORC1 activation. (A-D) Samples were obtained from the indicated strains as described in Fig 1B. *Left panels:* Transcript levels of the indicated ribosomal genes were measured at the indicated time points by quantitative real-time RT-PCR, normalized to the *act1* gene and expressed relative to those obtained at time 0’. Bars represent standard deviations calculated from two independent experiments with three technical replicates each. *Right panels:* Representative western blots showing phosphorylation of Rps6 at the indicated time points. α-Tubulin was used as loading control. Blots were performed with two biological replicates with similar results.

In a second approach, we generated a strain in which Gtr1 was locked in a constitutively active (GTP-bound) form (*gtr1^GTP^*) that was previously shown to activate TORC1 [17, 30]. A BlastP search using Gtr1 from *S. cerevisiae* (SGD:S000004590) as a bait detected a single *F. oxysporum* Gtr1 orthologue (FOXG_07552). ClustalW alignment [29] of the N-terminal region of Gtr1 from different organisms identified glutamine 86 (Q86) of *F. oxysporum* Gtr1 as the conserved residue to be replaced by leucine (L) in order to generate a GTP-locked Gtr1 variant [30–32] (S3 Fig). The *gtr1^Q86L^*allele was generated by site-directed mutagenesis and introduced in the wild-type strain by co-transformation with the *phleomycin B* resistance gene (S4 Fig). Analysis of several phleomycin-resistant co-transformants by PCR amplification and sequencing of the relevant genomic region identified a transformant carrying a discrete thymine (T) peak in position 257, indicating homologous replacement of the wild-type *gtr1* allele with *gtr1^Q86L^*in this strain (S4C Fig). In addition, we also obtained strains carrying both mutations by deleting the *tsc2* gene in the *gtr1^GTP^* background (S5 Fig). We noted that the TORC1 readouts in the *gtr1^GTP^*mutant were very similar to those of the wild-type strain upon a shift from Gln to −N, whereas those in the *gtr1^GTP^ tsc2*Δ double mutant were comparable to the *tsc2*Δ single mutant (Fig 2C and 2D). We thus conclude that inactivation of Tsc2 in *F. oxysporum* leads to constitutive activation of TORC1 while expression of *gtr1^GTP^* has no detectable effect on TORC1 activity under the conditions tested.

### Constitutive activation of TORC1 affects *Fusarium oxysporum* growth, development and stress response

Growth of the different fungal strains was analyzed in solid and liquid media. On plates, colony growth of the *tsc2*Δ mutant was generally reduced compared to that of the wild-type and the *tsc2*Δ^C^ strain, both in terms of diameter and hyphal density, but this effect was much more pronounced on minimal medium (Fig 3A). Importantly, the addition of rapamycin reduced colony growth to a similar extent in all strains tested. In liquid culture, biomass production of the *tsc2*Δ and *gtr1^GTP^ tsc2*Δ mutants was also significantly reduced compared to the wild-type strain, both in complete and minimal medium, but the effect was again more pronounced in the latter condition (Fig 3B). Moreover, microconidia production of the *tsc2*Δ and *gtr1^GTP^ tsc2*Δ mutants was significantly increased in complete, but not in minimal liquid medium (Fig 3C).

**Fig 3.**
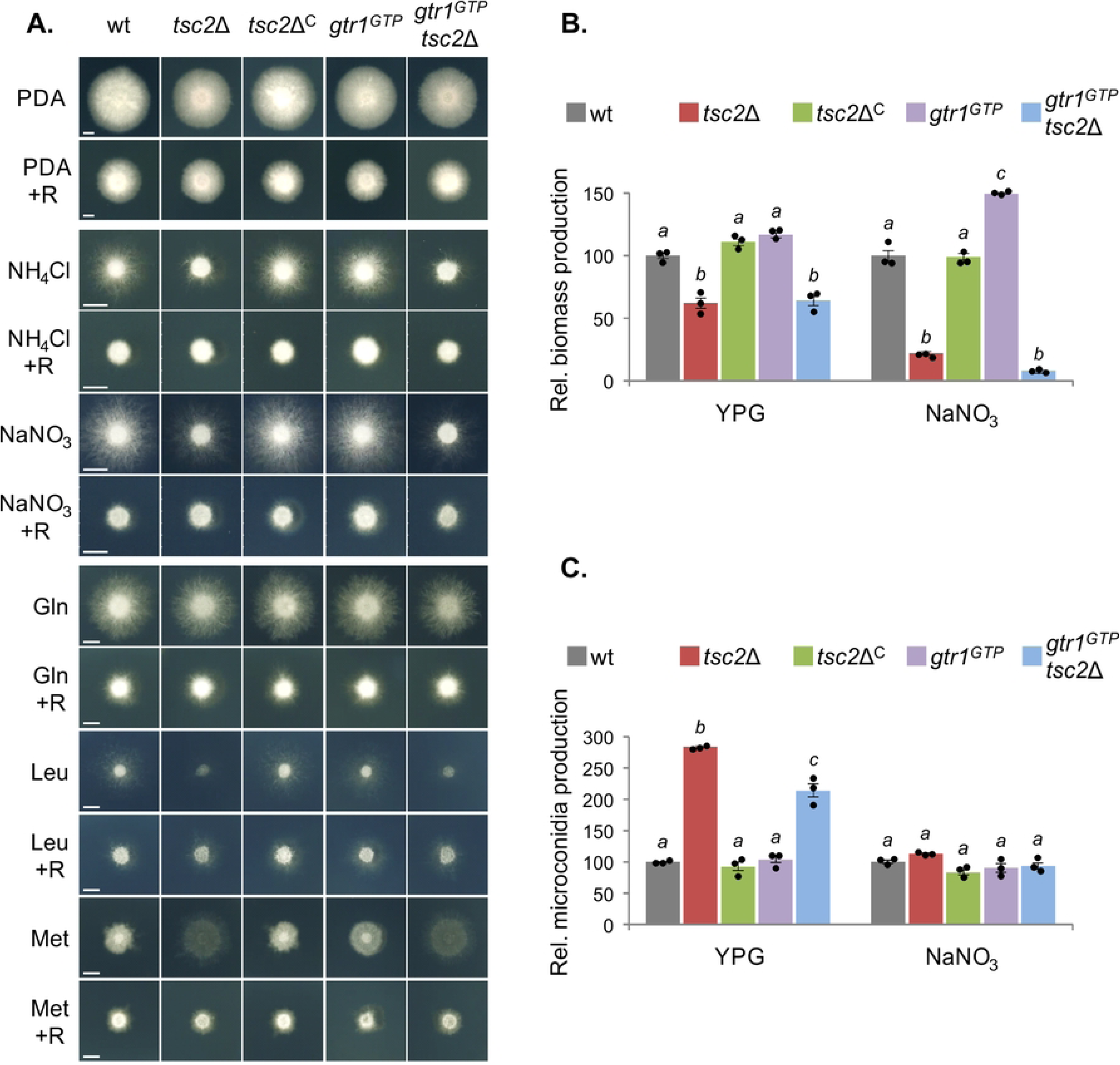
Constitutive activation of TORC1 affects growth and development of *Fusarium oxysporum.* (A) The indicated strains were spot-inoculated on Potato Dextrose Agar (PDA) or on minimal medium (MM) supplemented with the indicated inorganic or organic nitrogen source, with or without 2 ng ml^-1^ rapamycin (R). Cultures were grown at 28°C for 3-4 d and imaged. Scale bars = 5 mm. (B and C) Biomass (dry weight) (B) and microconidia production (C) of the indicated strains after 48 h growth in liquid YPG or in MM supplemented with NaNO_3_ shaking and static cultures, respectively. Values are represented relative to those of the wild-type strain. Values with the same letter are not significantly different (*p*<0.05) according to unpaired *t*-test. Bars represent standard deviations calculated from three independent experiments with two replicates each.

In contrast to the *tsc2*Δ mutant, growth of the *gtr1^GTP^* mutant was largely similar to that of the wild-type strain under most conditions tested, except for a slight reduction on PDA plates and an increase in biomass in liquid minimal medium (Fig 3A and 3B).

We next tested growth of the strains in the presence of different stresses and found that the *tsc2*Δ and *gtr1^GTP^ tsc2*Δ mutants displayed increased sensitivity to the cell wall targeting compound calcofluor white (CFW) and to high temperature stress (37°C) (Fig 4A and 4B). By contrast, no differences were detected in the presence of hyperosmotic (1 M NaCl, 1 M sorbitol) or oxidative (10 µg ml^-1^ menadione, 0.01% H_2_O_2_) stress conditions (not shown).

**Fig 4.**
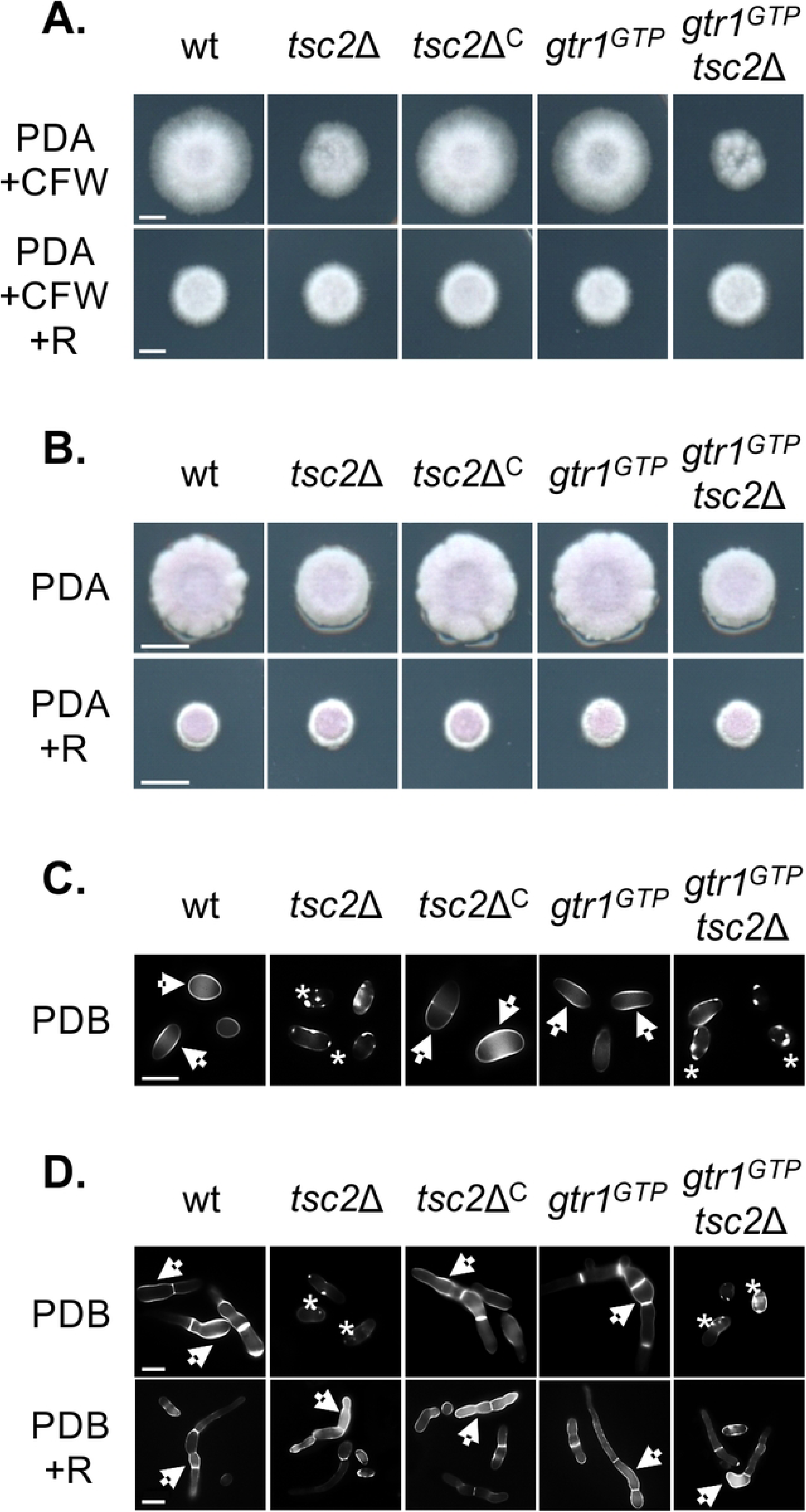
Constitutive activation of TORC1 increases sensitivity to cell wall and high temperature stress. (A and B) The indicated strains were spot-inoculated on PDA supplemented with 40 µg ml^-1^ calcofluor white (CFW) with or without 2 ng ml^-1^ rapamycin (R) and grown at 28°C for 3-4 d (A); or on PDA with or without 2 ng ml^-1^ rapamycin (R) and grown at 37°C for 4 d (B). Scale bars = 5 mm. (C and D) Microconidia of the indicated strains were germinated in PDB with and without 2 ng ml^-1^ rapamycin for 2 h (C) or 6-8 h (D) and stained with 6 mg ml^-1^ calcofluor white (CFW). Arrows highlight the uniform fluorescence signal along the cell wall while asterisks indicate irregular fluorescent punctae. Fluorescence microscopy images were taken at 100X magnification. Scale Bars = 5 µm.

To further investigate the integrity of the cell wall during conidial germination, fluorescence microscopy analysis was performed using the chitin-binding dye CFW. Two hours after inoculation in Potato Dextrose Broth (PDB), the microconidia of the wild-type, *tsc2*Δ^C^ and *gtr1^GTP^*strains showed a largely uniform fluorescence signal along the cell wall, whereas the *tsc2*Δ and *gtr1^GTP^ tsc2*Δ mutants showed a highly irregular distribution of fluorescence with a lower overall level, but strongly fluorescent punctae that were still detectable after 6-8 hours (Fig 4C and 4D). Moreover, the *tsc2*Δ and *gtr1^GTP^ tsc2*Δ mutants exhibited a marked delay in germination compared to the wild-type. Importantly, addition of rapamycin to the *tsc2*Δ and *gtr1^GTP^ tsc2*Δ mutants restored uniform distribution of the CFW fluorescence signal and germination to wild-type levels (Fig 4D). Taken together, these results indicate that constitutive activation of the TORC1 signaling pathway through *tsc2* deletion negatively affects *F. oxysporum* growth, development and response to different stress conditions.

### TORC1 activation blocks invasive hyphal growth of *Fusarium oxysporum*

Previous work established that fungal plant pathogenicity requires the correct activation of multiple virulence functions, including invasive hyphal growth, a process that is experimentally defined by the ability to penetrate across a cellophane membrane [7,33,34]. Here we found that, in contrast to the wild-type strain, the *tsc2*Δ and *gtr1^GTP^ tsc2*Δ mutants failed to penetrate cellophane membranes (Fig 5A). Importantly, invasive growth of these mutants was fully restored by addition of rapamycin (Fig 5B). In line with these findings, the *tsc2*Δ and *gtr1^GTP^ tsc2*Δ mutants caused less tissue maceration than the wild-type when inoculated on apple slices, indicative of reduced invasive growth on living fruit tissue (Fig 5C) [35]. Moreover, vegetative hyphal fusion, another virulence-related function that can be visualized by the formation of macroscopically visible hyphal aggregates [7,33,34], was also impaired in *tsc2*Δ and *gtr1^GTP^ tsc2*Δ, and to a lesser extent in *gtr1^GTP^*, and was fully rescued by rapamycin (Fig 5D). Collectively, these results demonstrate that TORC1 acts as a negatively regulator of virulence-related functions such as invasive hyphal growth.

**Fig 5.**
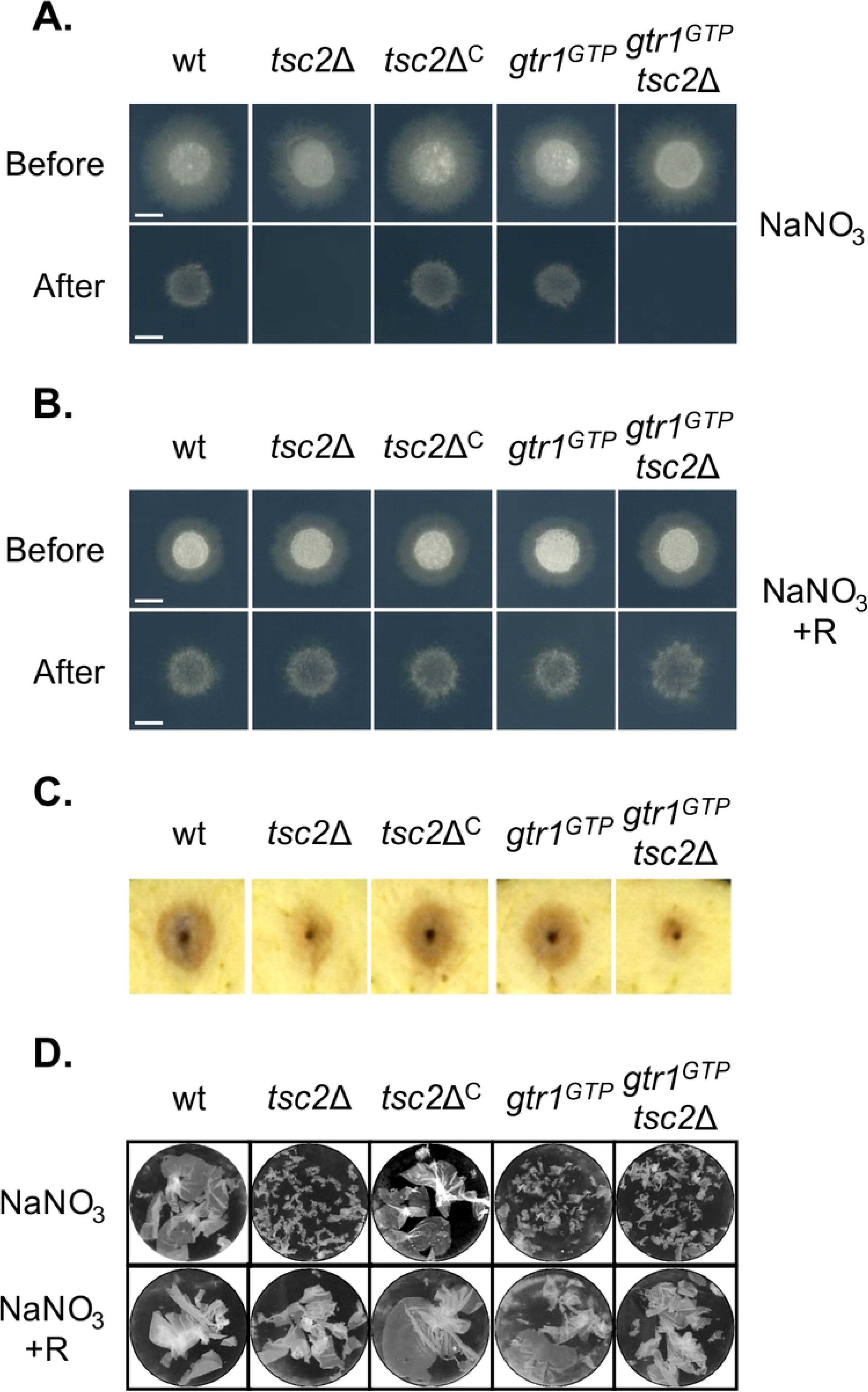
TORC1 negatively regulates virulence-related functions. (A and B) Cellophane penetration was determined on MM plates containing 50 mM NaNO_3_ without (A) or with (B) 2 ng ml^-1^ rapamycin (R). The indicated fungal strains were spot-inoculated and grown 4 d at 28°C on top of cellophane membranes (Before). The cellophane with the fungal colony was removed and plates were incubated for an additional day to determine the presence of mycelial growth on the plate, indicative of cellophane penetration (After). (C) Apple slices were spot-inoculated with the indicated strains and imaged after 3 days of incubation at 28°C and 100% humidity. (D) Hyphal aggregates forming 48 h after inoculation of the indicated strains in liquid MM supplemented with NaNO_3_, with or without 2 ng ml^-1^ rapamycin (R). Cultures were vortexed to dissociate weakly adhered hyphae and images were taken with a binocular microscope.

### Regulation of TORC1 is essential for virulence in *Fusarium oxysporum*

We next asked whether correct modulation of TORC1 signaling is important for the ability of *F. oxysporum* to infect tomato plants. Tomato seedlings, whose roots were inoculated with microconidia of the wild-type or the complemented *tsc2*Δ^C^ strain developed progressive wilt symptoms and were mostly dead around day 25 post-inoculation (dpi) (Figs 6A and S6). On the contrary, plants inoculated with *tsc2*Δ and *gtr1^GTP^ tsc2*Δ displayed much milder symptoms and a markedly reduced mortality rate. Furthermore, fungal burden in roots and stems at 10 dpi was significantly lower in plants inoculated with *tsc2*Δ and *gtr1^GTP^tsc2*Δ compared to plants inoculated with the wild-type or the *tsc2*Δ^C^ strain. Interestingly, plants inoculated with *gtr1^GTP^*also exhibited a reduction in mortality. However, in contrast to plants inoculated with *tsc2*Δ and *gtr1^GTP^ tsc2*Δ, those inoculated with *gtr1^GTP^* contained more fungal biomass than those infected with the wild-type strain (Fig 6A and 6B).

**Fig 6.**
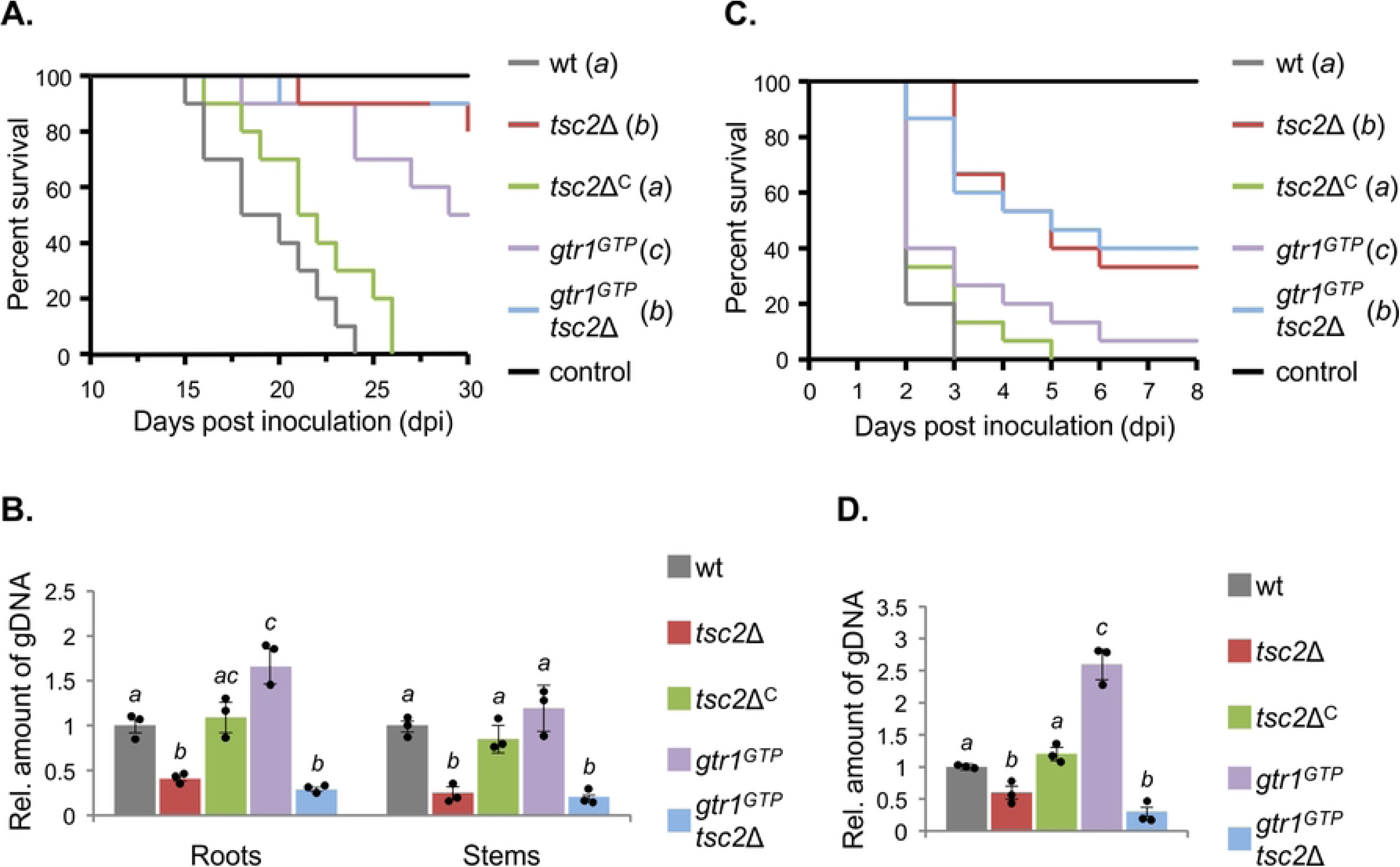
TORC1 regulation is important for virulence of *Fusarium oxysporum* on plant and animal hosts. (A and C) Kaplan-Meier plots showing the survival of tomato plants (A) or *Galleria mellonella* larvae (C) inoculated with the indicated fungal strains. Number of independent experiments = 3; 10 plants or 15 larvae per treatment. Data shown are from one representative experiment. Treatments with the same letter are not significantly different (*P* <0.05) according to log-rank test. (B and D) Quantitative real-time PCR was used to measure the relative amount of fungal DNA in total genomic DNA extracted from tomato roots and stems 10 days after inoculation (B) or from *G. mellonella* larvae 2 days after inoculation (D). Fungal burden is expressed relative to that measured in plants or larvae infected with the wild-type strain. Values with the same letter are not significantly different (*p*<0.05) according to unpaired *t*-test. Bars represent standard deviations calculated from three independent experiments with two replicates each.

Because *F. oxysporum* can cause disseminated infections in humans [3], we tested the virulence of the different strains in larvae of the wax moth *G. mellonella*, an invertebrate model that is widely used to study microbial pathogens of humans including *F. oxysporum* [36]. Inoculation with microconidia of the wild-type or the complemented *tsc2*Δ^C^ strain resulted in killing of all larvae at 3-5 dpi (Fig 6C). However, animals inoculated with *tsc2*Δ and *gtr1^GTP^ tsc2*Δ, and to a lesser extent those inoculated with *gtr1^GTP^*, showed significantly lower mortality rates. Moreover, similar to the plant infection assays, fungal burden at 2 dpi was lower in *G. mellonella* larvae infected with the *tsc2*Δ and *gtr1^GTP^ tsc2*Δ mutants and higher in those infected with *gtr1^GTP^*, as compared to larvae infected with the wild-type and the *tsc2*Δ^C^ strain (Fig 6D). Altogether, these results establish an important function of correct regulation of TORC1 during *F. oxysporum* infection on plant and animal hosts.

## DISCUSSION

During the infection process, fungal phytopathogens sense an array of environmental signals and must generate the appropriate developmental and morphogenetic responses to ensure successful colonization of the plant host [37]. It has been proposed that nitrogen limitation acts as a key signal in plant pathogens to trigger the expression of virulence genes [7, 38]. Because TORC1 promotes cell growth and proliferation in response to nutrient sufficiency [39], we hypothesized that uncontrolled activation of the TORC1 pathway should negatively impact virulence in *F. oxysporum*. Indeed, previous work in *M. oryzae* showed that mutations affecting carbon and nitrogen metabolism, which cause inappropriate activation of the TORC1 pathway, also impair appressorium formation [22, 40].

We attempted constitutive activation of the TORC1 signaling pathway in *F. oxysporum* by combining two different genetic approaches: (1) inactivation of the negative regulator Tsc2 resulting in overstimulation of the positive TORC1 regulator Rheb [41, 42], and (2) expression of a GTP-locked allele (*gtr1^GTP^*) of the positive TORC1 regulator Gtr1 [43]. We found that the first strategy indeed resulted in nutrient status-independent activation of TORC1 signaling, based on transcription of ribosomal genes and phosphorylation of Rps6, two readouts commonly used to measure TORC1 activation status [24–27]. By contrast, expression of *gtr1^GTP^* in *F. oxysporum* had no detectable effect on TORC1 activation. A possible explanation is provided by the previous finding that expression of *gtr1^GTP^* in *S. cerevisiae* only leads to partial activation of TORC1 [44]. Moreover, recent work in *S. pombe* suggests that the role of Rag GTPases is more complex than previously recognized and also involves attenuation, rather than activation, of TORC1 [45–47]. Our data suggest that deletion of *tsc2* is the most effective approach for constitutive activation of TORC1 in *F. oxysporum*.

### Constitutive activation of TORC1 alters growth and stress response of *Fusarium oxysporum*

To adapt growth to different environmental situations, fungi exquisitely regulate nutrient uptake and utilization. Under nutrient limiting conditions, TORC1 is turned off and the production of transport proteins for nitrogenous compounds increases [13, 48]. On the contrary, under optimal growth conditions transporters and permeases in the plasma membrane are downregulated in a TORC1 dependent manner via ubiquitination and endocytosis [49]. Here we propose that reduced growth of the *tsc2*Δ mutants on non-preferred nitrogen sources is a consequence of inappropriate TORC1 activation. This is supported by the finding that the growth differences between the *tsc2*Δ strains and the wild-type were levelled when TORC1 was inhibited by rapamycin.

In addition to affecting growth, inappropriate activation of TORC1 also increased sensitivity of *F. oxysporum* to cell wall and high temperature stresses, and this effect was reversed by rapamycin. The fungal cell wall provides a rigid cellular border and represents the first line of defense against adverse environmental conditions. In *F. oxysporum*, the cell wall is mainly composed of glucans, chitin and glycoproteins [50] and its integrity is essential for development and virulence [51–53]. Interestingly, *S. cerevisiae* cells expressing a hyperactive *TOR1* allele were impaired in β-1-3-glucan synthesis and exhibited increased sensitivity to cell wall perturbing compounds [54]. Furthermore, in *F. graminearum* loss of Sit4, a phosphatase that is inactivated by TORC1, caused increased sensitivity to cell wall damaging agents and to high temperature stress [21]. Interestingly, *F. oxysporum* mutants in the cell wall integrity (CWI) mitogen activated protein kinase (MAPK) cascade also display increased sensitivity to both cell wall and high temperature stress [55]. Although the functional link between CWI and TORC1 signaling pathways is not fully understood, a crosstalk between these two pathways has previously been suggested [21,54,56,57].

Treatment of *F. oxysporum* germlings with the chitin-binding dye CFW produced intense staining of the cell wall and septae as previously reported [51], whereas in the *tsc2*Δ strains an abnormal punctated pattern of chitin deposition was observed. This was concomitant with a marked delay in germ tube growth. Abnormal chitin aggregates have also been observed in *S. cerevisiae* cells expressing a hyperactive *TOR1* allele [54]. Because correct recycling of the chitin synthase Chs3 is required for polarized hyphal growth [58], we speculate that constitutive activation of TORC1 could interfere with this process, as shown for other proteins localized in the plasma membrane [49], thereby affecting cell wall biogenesis and germination. Importantly, chemical inhibition of TORC1 fully restored wild-type cell wall biogenesis and germination in the *tsc2*Δ mutants. We further found that microconidia production is increased in the *tsc2*Δ strains while biomass production is reduced. Previously, inactivation of the protein phosphatases Sit4 or Ppg1, both of which are inactive when TORC1 signaling is on, was also shown to increase conidiation and reduce mycelial growth in *F. graminearum* [21].

### TORC1 inhibits fungal virulence functions

Cellophane penetration is routinely used as a measure of invasive hyphal growth in *F. oxysporum*, due to its similarities with the agar invasion assay in yeast [59] and because it highly correlates with the virulence phenotype on tomato plants [7,33,60]. Cellophane penetration, as well as other virulence-related processes such as vegetative hyphal fusion or invasive growth on apple slices, were markedly inhibited in the *tsc2*Δ strains, while rapamycin-mediated TORC1 inactivation restored these functions. Together with previous evidence [7,34,61], our findings suggest a broadly conserved role of TORC1 as a negative regulator of infection-related processes in plant and human fungal pathogens. The finding that *F. oxysporum* mutants lacking *tsc2* are significantly reduced in their ability to invade and kill tomato plants or the non-vertebrate animal model *G. mellonella*, further highlights the relevance of the TORC1 pathway in fungal pathogenicity. The fact that inactivation of Tsc2 attenuates virulence of *F. oxysporum* on both plants and animals suggests that TORC1- regulated mechanisms are relevant for infection in both types of hosts. Previous studies established that both the invasive growth MAPK Fmk1 and the CWI MAPK Mpk1 control critical steps during the infection process of *F. oxysporum* [5,8,53,55,62]. Collectively, these findings suggest that the virulence phenotypes observed upon TORC1 deregulation might be, at least in part, associated with defects in MAPK signaling. In line with this, inactivation of the protein phosphatase Sit4, which is negatively regulated by TORC1, increases cell wall stress and reduces virulence in *F. graminearum* [21]. Both Sit4 and another phosphatase called Ppg1 interact with protein phosphatase Msg5, a negative regulator of the CWI MAPK pathway [21]. Importantly, Msg5 was recently shown to regulate cell wall integrity and virulence in *F. oxysporum* [53]. Somewhat unexpectedly, expression of a *gtr1^GTP^*allele, which was previously shown to activate TORC1 in different organisms [17, 30], was ineffective in *F. oxysporum* since no constitutive TORC1 activation was detected under the experimental conditions used in this study. Interestingly, however, the *gtr1^GTP^* mutant displayed some striking phenotypes such as increased proliferation as determined by higher fungal biomass in liquid cultures and under infection conditions, as well as attenuated virulence on plant and animal hosts. The discrepancy in phenotypes between the *tsc2*Δ and the *gtr1^GTP^* mutants in most assays carried out in this study (growth, stress, development, conidiation, invasive growth) suggest that the mechanisms underlying the virulence defects of these two mutants could be different. Importantly, inactivation of Tsc2 in a *gtr1^GTP^* background recapitulated all the phenotypes of the *tsc2*Δ single mutant and no intermediate phenotypes or additive effects were detected, suggesting that targeted deletion of *tsc2* is epistatic to *gtr1^GTP^* and that both components function in the same pathway.

In summary, our results support a conserved role of TORC1 as a negative regulator of fungal virulence. Moreover, the finding that fungal infection can be targeted by inappropriate TORC1 activation opens new avenues for antifungal discovery.

## MATERIALS AND METHODS

### Fungal isolates and culture conditions

*Fusarium oxysporum* f. sp. *lycopersici* race 2 isolate Fol4287 (FGSC 9935) and the derived isogenic mutants were stored as microconidial suspensions at - 80°C in 30% glycerol. For the extraction of genomic DNA and microconidia production, cultures were grown in Potato Dextrose Broth (PDB) at 28°C [63]. For the determination of colony growth, 2 x 10^4^ microconidia were spotted onto Potato Dextrose Agar (PDA), minimal medium [64] with 25 mM of the indicated nitrogen source, or Yeast extract Peptone Glucose Agar (YPGA). When needed, media were supplemented with the indicated concentrations of rapamycin (R) or calcofluor white (CFW) (both from Sigma-Aldrich, Madrid, Spain). Cultures were incubated at the indicated temperatures for the specified time periods. Conidiation was quantified in Yeast extract Peptone Glucose (YPG) or in 25 mM sodium nitrate minimal medium (NaNO_3_) static liquid cultures grown as described [65, 66]. For fungal biomass quantification, 2.5 x 10^6^ microconidia ml^-1^ were inoculated in YPG or NaNO_3_ and cultures were maintained under standard conditions (28°C and 170 rpm). After 48 h the mycelium was harvested, dried, and weighed. Invasion assays on cellophane membranes (Colorless, Manipulados Margok, Gipuzkoa, Spain) were performed as described [7, 33] using solid minimal medium supplemented with 50 mM NaNO_3_ with or without 2 ng ml^-1^ rapamycin. For macroscopic analysis of hyphal fusion and agglutination, fungal strains were grown 48 h in minimal medium supplemented with 50 mM NaNO_3_ with or without 2 ng ml^-1^ rapamycin. Cultures were vortexed to dissociate weakly adhered hyphae and aliquots were transferred to 12-well cell culture plates and imaged using a Leica DFC 300 FX digital camera coupled to a Leica binocular microscope driven by Leica IM50 4.0 software (Leica, Wetzlar, Germany). ImageJ software (National Institutes of Health, Bethesda, MD, USA) was used for contrast adjustment. For gene expression analysis, freshly obtained microconidia were germinated 16 h in PDB. Germlings were harvested by filtration, washed three times in sterile water and transferred: 1) to nitrogen-free minimal medium (−N) for 1 h before shifted to −N or glutamine minimal medium (Gln) for an additional hour; 2) to Gln for 1 h before shifted to −N or −N + 100 ng ml^-1^ rapamycin (−N+R) for an additional hour; or 3) to Gln for 1 h before shifted to Gln or Gln + 100 ng ml^-1^ rapamycin (Gln+R) for an additional hour. All experiments included three replicates and were performed at least twice with similar results.

### Generation of mutant strains

PCR reactions were routinely performed with VELOCITY DNA Polymerase (Bioline, London, UK) using an MJ Mini Personal Thermal Cycler (Bio-Rad, Madrid, Spain). All fungal transformations and purification of the transformants by monoconidial isolation were performed as described previously [63]. Targeted replacement of the entire coding region of the *F. oxysporum tsc2* gene was performed as depicted (S2 Fig) using the split-marker method. Plasmid pAN7-1, containing the *hygromycin B* resistance gene (*hyg*) under the control of the *Aspergillus nidulans gpdA* promoter and *trpC* terminator [67] was used. Transformants were genotyped by PCR (not shown) and Southern blot analysis and the *tsc2*Δ*#6* mutant was used for further experiments and complementation (S2 Fig). Complementation of *tsc2*Δ with a PCR fragment encompassing the *tsc2* wild-type allele was done by co-transformation with the *phleomycin B* resistance gene under the control of the *A. nidulans gpdA* promoter and *trpC* terminator amplified from plasmid pAN8-1. Several phleomycin-resistant co-transformants were analyzed for the presence of a functional *tsc2* gene and, among them, we specially selected those who had lost hygromycin resistance, therefore with *tsc2* integrated at its original locus. Co-transformants were genotyped by PCR (not shown) and Southern blot analysis, and *tsc2*Δ^C^*#1* was selected for further experiments (S2 Fig).

For the generation of a strain in which Gtr1 is expressed in a constitutively active (GTP-bound) form (*gtr1^GTP^*), a *gtr1^Q86L^* allele was constructed by site-directed mutagenesis (S4 Fig). Briefly, specific primers harboring the *gtr1^256CTA258^* mutation (S4 Fig) were used to amplify two fragments which were subsequently assembled by a fusion PCR method [68]. The obtained DNA fragment containing the *gtr1^Q86L^* allele was cloned into the pGEM-T vector (Promega, Madison, WI, USA), verified by Sanger sequencing and used to co-transform protoplasts of the *F. oxysporum* wild-type strain together with the *phleomycin B* resistance gene. Several phleomycin-resistant transformants were analyzed by amplification and sequencing of the relevant *gtr1* fragment. The fluorogram of *gtr1^GTP^#1*, showing a discrete thymine (T) peak in position 257, indicates the sole presence of the *gtr1^Q86L^* allele in this strain (S4C Fig). The expression of exclusively the *gtr1^Q86L^* allele was further confirmed by RT-PCR and sequencing (not shown). Targeted replacement of *tsc2* in a *gtr1^GTP^* background was performed as described above using the split-marker method. Transformants were genotyped by PCR (not shown) and Southern blot analysis (S5 Fig).

### Nucleic acid manipulations and real time quantitative RT-PCR

Total RNA and gDNA were extracted from *F. oxysporum* mycelia following previously reported protocols [69, 70]. Quality and quantity of the extracted nucleic acids were determined by running aliquots in ethidium bromide-stained agarose gels and by spectrophotometric analysis in a NanoDrop ND-1000 spectrophotometer (NanoDrop Technologies, Wilmington, DE, USA). Routine nucleic acid manipulations were performed according to standard protocols [71]. Quantitative RT-PCR was performed as described previously [7, 72] using FastStart Essential DNA Green Master (Roche Diagnostics SL, Barcelona, Spain) in a CFX Connect Real-Time System (Bio-Rad, Madrid, Spain). Transcript levels were calculated by comparative ΔCt and normalized to *act1*.

### Western blot

Proteins were extracted using a reported procedure [73, 74] involving solubilization from lyophilized mycelial biomass with NaOH, followed by precipitation with trichloroacetic acid (TCA). Aliquots were resolved in 10-12% SDS-polycrylamide gels and transferred to nitrocellulose membranes with a Trans-Blot Turbo Transfer System (Bio-Rad, Madrid, Spain) for blotting. Western blots were reacted with Phospho-(Ser/Thr) Akt Substrate Antibody (1:10000, #9611 Cell Signaling Technology, Danvers, MA, USA) as primary and with Anti-Rabbit IgG-Peroxidase Antibody (1:10000; A1949 Sigma-Aldrich, Madrid, Spain) as secondary, or with Anti-α-Tubulin Antibody (1:10000; T6119 Sigma-Aldrich, Madrid, Spain) as primary and Anti-Mouse IgG-Peroxidase Antibody (1:10000; A4416 Sigma-Aldrich, Madrid, Spain) as secondary. Proteins were detected with ECL (Amersham Biosciences, Madrid, Spain).

### Fluorescence microscopy

*F. oxysporum* microconidia were germinated in PDB with or without 2 ng ml^-1^ rapamycin for 2h or 6-8 h and stained with 6 mg ml^-1^ CFW. Images were acquired at 100X magnification using a Leica DFC 300 FX digital camera coupled to a Leica DMR microscope driven by Leica IM50 4.0 software (Leica, Wetzlar, Germany). ImageJ software (National Institutes of Health, Bethesda, MD, USA) was used for contrast adjustment.

### Infection assays

Tomato root inoculation assays were performed as described [63] using 2- week-old tomato seedlings, cultivar Monika (Syngenta, Almería, Spain). Severity of disease symptoms and plant survival were recorded daily for 30 dpi. 10 plants were used for each treatment. Data were analyzed with GraphPad Prism 5 software. Quantification of fungal biomass *in planta* was performed as described [75] using total gDNA extracted from tomato roots or stems infected with *F. oxysporum* strains at 10 dpi. Relative amounts of fungal gDNA were calculated by comparative ΔCt of the *F. oxysporum act1* gene normalized to the tomato *EFα1* gene. Infection assays on apple slices, cultivar Golden Delicious, were performed as described [35]. Colonization and maceration of the fruit tissue were monitored daily for 3-5 dpi. *Galleria mellonella* infection assays were performed as described [36]. *G. mellonella* larvae (Nutri-reptil, Córdoba, Spain) were maintained in plastic boxes for 2-3 d before the infection. Fifteen larvae were used for each treatment. A Burkard Auto Microapplicator (0.1-10 µl; Burkard Manufacturing Co. Limited, Hertfordshire, UK) with a 1 ml syringe (Terumo Medical Corporation, Somerset, NJ, USA) was used to inject 8 µl of a 1.6 x 10^5^ microconidial suspension into the hemocoel of each larva. After injection, larvae were incubated in glass containers at 30°C. Survival was recorded daily for 10 dpi. Data were analyzed with GraphPad Prism 5 software. Quantification of fungal biomass in *G. mellonella* larvae was performed as described using total gDNA extracted from animals infected with *F. oxysporum* strains at 2 dpi. Relative amounts of fungal gDNA were calculated by comparative ΔCt of the *F. oxysporum act1* gene normalized to the *G. mellonella gallerimycin* gene. Virulence experiments were performed at least three times with comparable results.

## ACKNOWLEDGEMENTS

We are grateful to Esther Martínez Aguilera for valuable technical assistance and to Syngenta (Spain) for kindly supplying tomato seeds of cultivar Monika. G.Y.V.N. was funded by a PhD fellowship from IFARHU, Panama. This work was supported by grants PID2019-108045RB-I00 from the Spanish Ministerio de Ciencia e Innovación and P20_00179 from Junta de Andalucía to A.D.P.

## SUPPORTING INFORMATION CAPTIONS

**S1 Fig. Amino acid sequence alignment of the Tsc2 GTPase-activating domain.**

The predicted *F*. *oxysporum tsc2* gene product (FOXG_01632) was aligned with the Tsc2 GTPase-activating domain from *Schizosaccharomyces pombe*, *Homo sapiens* and *Neurospora crassa*. Conserved and similar residues are shaded in black and grey, respectively.

**S2 Fig. *Fusarium oxysporum tsc2* knockout strategy.**

(A) *F. oxysporum tsc2* locus and targeted gene disruption construct. (B) Southern blot analysis for knockout identification. gDNA of the indicated strains was treated with *BamH*I, separated on a 0.7% agarose gel, transferred to a nylon membrane and hybridized with the DNA probe indicated in (A). (C) Southern blot analysis for complemented strain identification. gDNA of the indicated strains was treated with *BamH*I, separated on a 0.7% agarose gel, transferred to a nylon membrane and hybridized with the DNA probe indicated in (A).

**S3 Fig. Amino acid sequence alignment of the N-terminal region of Gtr1.** The predicted *F*. *oxysporum gtr1* gene product (FOXG_07552) was aligned with the N-terminal region of Gtr1 from *Saccharomyces cerevisiae*, *Schizosaccharomyces pombe*, *Homo sapiens* and *Neurospora crassa*. Conserved and similar residues are shaded in black and grey, respectively. The conserved glutamine (Q) replaced by leucine (L) to generate the Gtr1^GTP^ form is boxed in red.

**S4 Fig. Generation of *Fusarium oxysporum gtr1^GTP^*by site-directed mutagenesis.**

(A) *F. oxysporum gtr1* locus and site-directed mutagenesis representation. (B-D) Analysis of the wild-type (B) and of two different co-transformants (C and D) by PCR amplification of gDNA and subsequent sequencing of a fragment encompassing the CAA to CTA substitution. Original fluorograms are shown indicating the nucleotides mutated in each strain.

**S5 Fig. Generation of a *Fusarium oxysporum gtr1^GTP^ tsc2*Δ strain.**

(A) *F. oxysporum gtr1* locus and site-directed mutagenesis representation (S4A Fig) and *tsc2* locus and targeted gene disruption construct (S2A Fig). (B) Southern blot analysis for *tsc2* knockout identification. gDNA of the indicated strains was treated with *BamH*I, separated on a 0.7% agarose gel, transferred to a nylon membrane and hybridized with the DNA probe indicated in (A).

**S6 Fig. Incidence of *Fusarium* wilt on tomato plants.**

Groups of 10 tomato plants (cv Monika) were dip-inoculated with a microconidial suspension of the indicated fungal strains. The severity of disease symptoms was monitored periodically using an index ranging from 1 (healthy plant) to 5 (dead plant) (see image below). Bars represent standard errors calculated from 10 plants.

## Notes

### Competing Interest Statement

The authors have declared no competing interest.

